# Electrophysiological network alterations in adults with copy number variants associated with high neurodevelopmental risk

**DOI:** 10.1101/753145

**Authors:** Diana C. Dima, Rachael Adams, Stefanie C. Linden, Alister Baird, Jacqueline Smith, Sonya Foley, Gavin Perry, Bethany C. Routley, Lorenzo Magazzini, Mark Drakesmith, Nigel Williams, Joanne Doherty, Marianne B.M. van den Bree, Michael J. Owen, Jeremy Hall, David E. J. Linden, Krish D. Singh

**Affiliations:** Cardiff University Brain Research Imaging Centre (CUBRIC), School of Psychology, Cardiff University, Maindy Road, Cardiff, CF24 4HQ, United Kingdom; Neuroscience and Mental Health Research Institute (NMHRI), Cardiff University, Maindy Road, Cardiff, CF24 4HQ, United Kingdom; Division of Psychological Medicine and Clinical Neurosciences, School of Medicine, Cardiff University, Cardiff, CF24 4HQ, UK; MRC Centre for Neuropsychiatric Genetics and Genomics, School of Medicine, Cardiff University, Maindy Road, Cardiff, CF24 4HQ, United Kingdom; School of Mental Health and Neuroscience, Faculty of Health, Medicine and Life Sciences, Maastricht University, Maastricht, The Netherlands

## Abstract

Rare copy number variants associated with increased risk for neurodevelopmental and psychiatric disorders (referred to as ND-CNVs) are characterized by heterogeneous phenotypes thought to share a considerable degree of overlap. Altered neural integration has often been linked to psychopathology and is a candidate marker for potential convergent mechanisms through which ND-CNVs modify risk; however, the rarity of ND-CNVs means that few studies have assessed their neural correlates. Here, we used magnetoencephalography (MEG) to investigate resting-state oscillatory connectivity in a cohort of 42 adults with ND-CNVs, including deletions or duplications at 22q11.2, 15q11.2, 15q13.3, 16p11.2, 17q12, 1q21.1, 3q29, and 2p16.3, and 42 controls. We observed decreased connectivity between occipital, temporal and parietal areas in participants with ND-CNVs. This pattern was common across genotypes and not exclusively characteristic of 22q11.2 deletions, which were present in a third of our cohort. Furthermore, a data-driven graph theory framework enabled us to successfully distinguish participants with ND-CNVs from unaffected controls using differences in node centrality and network segregation. Together, our results point to alterations in electrophysiological connectivity as a putative common mechanism through which genetic factors confer increased risk for neurodevelopmental and psychiatric disorders.

## Introduction

A number of rare genetic variants occurring through the deletion or duplication of chromosomal segments are associated with significantly increased risk for a range of neurodevelopmental disorders (ND), including schizophrenia, autism spectrum disorder (ASD), and developmental delay (Kirov et al., 2014). Although the underlying mechanisms remain poorly understood, these copy number variants (referred to hereafter as ND-CNVs) are thought to increase the risk for psychopathology through alterations in neural structure and function. Thus, neuroimaging studies in participants with ND-CNVs provide a unique opportunity to study intermediate phenotypes of mental disorders.

Recent work suggests that CNV-specific phenotypic outcomes are limited, pointing instead to a large degree of similarity across phenotypes associated with different ND-CNVs (Chawner et al., 2019; Niarchou et al., 2019). Focusing on convergent neural alterations across different genotypes can thus help elucidate the mechanisms linking ND-CNVs at different loci to a shared psychopathology and increase in neurodevelopmental risk.

Failures of functional neural integration are considered a hallmark of schizophrenia (Friston et al., 2016; Dong et al., 2018) and have also been found in ASD (Hull et al., 2017; O’Reilly et al., 2017) and across diagnostic boundaries (Sha et al., 2019). ND-CNVs are thought to increase disorder risk by acting on large-scale neural integration through molecular and cellular mechanisms (Karayiorgou et al., 2010). Synchronous oscillatory activity thought to support communication between brain areas is thus of particular interest as a potential biomarker of neurodevelopmental risk, and can be measured at rest using electro- and magneto-encephalography (EEG/MEG).

However, the rarity of ND-CNVs means that evidence of their functional connectivity correlates is scarce. Of the genetic imaging studies conducted so far, most have focused on the 22q11.2 deletion syndrome. This deletion is associated with a number of physical phenotype manifestations as well as high risk for psychopathology (Jonas et al., 2014; Niarchou et al., 2014; Schneider et al., 2014; Owen and Doherty, 2016). The presence of a 22q11.2 deletion has been linked to alterations in brain structure and function (Zinkstok and Amelsvoort, 2005; Boot and van Amelsvoort, 2013; Reddaway et al., 2018; Sun et al., 2018), including disrupted structural connectivity (Ottet et al., 2013; Villalón-Reina et al., 2019). Although fewer studies have investigated functional connectivity, they report similarly disrupted networks using functional MRI (Debbané et al., 2012; Padula et al., 2015; Scariati et al., 2016b) and EEG (Tomescu et al., 2014).

Despite emerging evidence of white matter alterations associated with other ND-CNVs (Berman et al., 2016; Drakesmith et al., 2019; Silva et al., 2019), very few studies have investigated their functional correlates. Recent electrophysiological research reported increased beta-band activity in participants with 15q11.2-q13.1 duplications (Frohlich et al., 2016, 2019) and 16p11.2 deletions (Hinkley et al., 2019), as well as delayed evoked responses in the latter (Berman et al., 2016; Jenkins et al., 2016). Based on current evidence it is difficult to assess the extent of functional connectivity alterations, especially for rarer ND-CNVs.

To address this, we investigated oscillatory connectivity measured with MEG in participants with ND-CNVs at nine different loci. Given the common phenotypic outcomes associated with ND-CNVs (Chawner et al., 2019), this approach can identify convergent endophenotypes of potentially higher clinical relevance. Because a third of our cohort had 22q11.2 deletions, we also investigated alterations in connectivity separately in this subgroup and in the group of participants with other ND-CNVs. This allowed us to assess the specificity of the effects, especially considering previous findings of widespread neural alterations associated with 22q11.2 deletions.

In both subgroups, we found evidence of disrupted alpha and beta-band oscillatory connectivity in posterior brain regions. Furthermore, using graph theory measures of network topology and information transfer, we were able to identify participants with ND-CNVs based on their individual connectivity maps. The two approaches highlighted common patterns of dysconnectivity in participants with ND-CNVs, as well as specific network features that might be linked to CNV pathogenicity.

## Methods

### Participants

MEG data were acquired in 42 adults with ND-CNVs targeted for their high penetrance for neurodevelopmental disorders (22 female; mean age 38.5±12.5 years). ND-CNVs at nine different loci were represented in the cohort, with 14 (33%) participants carrying 22q11.2 deletions (Table 1).

**Table 1.** Participant information (ND-CNV status, demographic information, number of psychiatric diagnoses, and IQ). Note that the 22q11.2 deletion group included two atypical deletions and one adjacent deletion.

| CNV and critical region (hg19) | Total N | N female | Age (mean±SD) | N diag (mean±SD) | VIQ (mean±SD) | PIQ (mean±SD) | FSIQ (mean±SD) |
| --- | --- | --- | --- | --- | --- | --- | --- |
| *All CNV* | 42 | 22 | 38.53±12.55 | 2.93±2.54 | 88.72±18.84 | 96.63±14.63 | 91.24±17.15 |
| 22q11.2del<br>chr22:19,037,332-21,466,726 | 14 | 9 | 38.97±16.43 | 3±2.07 | 86.28±21.23 | 95.28±15.6 | 89.93±17.88 |
| 22q11.2dup<br>chr22:19,037,332-21,466,726 | 4 | 13 | 38.32±10.44 | 2.89±2.78 | 90±17.77 | 97.33±14.36 | 91.89±17.08 |
| 17q12dup<br>chr17:34,815,904-36,217,432 | 2 |  |  |  |  |  |  |
| 16p11.2del<br>chr16:29,650,840-30,200,773 | 1 |  |  |  |  |  |  |
| 16p11.2dup (distal)<br>chr16:28,823,196-29,046,783 | 1 |  |  |  |  |  |  |
| 15q13.1-13.3del (BP4-5)<br>chr15:31,080,645-32,462,776 | 2 |  |  |  |  |  |  |
| 15q13.1-13.3dup (BP4-5)<br>chr15:31,080,645-32,462,776 | 1 |  |  |  |  |  |  |
| 15q11.2del (BP1-2)<br>chr15:22,805,313-23,094,530 | 7 |  |  |  |  |  |  |
| 15q11.2dup (BP1-2)<br>chr15:22,805,313-23,094,530 | 1 |  |  |  |  |  |  |
| 15q11.2q12dup (BP2-3; PWS/AS)<br>chr15:22,805,313-28390339 | 1 |  |  |  |  |  |  |
| 3q29del<br>chr3:195,720,167-197,354,826 | 1 |  |  |  |  |  |  |
| 2p16.3del (NRXN1)<br>chr2:50145643-51259674 | 1 |  |  |  |  |  |  |
| 1q21.1del<br>chr1:146,527,987-147,394,444 | 5 |  |  |  |  |  |  |
| 1q21.1dup<br>chr1:146,527,987-147,394,444 | 1 |  |  |  |  |  |  |
| *Controls* | 42 | 22 | 33.35±9.58 | - | - | - | - |

Recruitment was performed through NHS genetics clinics and relevant support groups within the UK. Written consent was obtained in accordance with The Code of Ethics of the World Medical Association (Declaration of Helsinki), and all procedures were approved by the South East Wales Research Ethics Committee.

All participants with ND-CNVs were assessed using the Psychiatric Assessment Schedule for Adults with Developmental Disability (PAS-ADD; Moss et al., 1997). 24 were also assessed using the Structured Interview for Prodromal Symptoms (SIPS; Miller et al., 2003), and 26 using the Structured Clinical Interview for DSM (SCID II; First and Gibbon, 2004). Diagnoses were assigned by research psychologists and verified by an adult psychiatrist using the Diagnostic and Statistical Manual for Mental Disorders, 4^th^ and 5^th^ Editions. Of the 42 participants, 34 had at least one diagnosis; 22 had anxiety disorders, 15 had mood disorders, and 15 had neurodevelopmental disorders (including 6 with intellectual disability and 6 with autism spectrum disorders). Five participants exhibited full or attenuated psychotic symptoms, of whom four met the criteria for a psychotic disorder, with two schizophrenia diagnoses in the 22q11.2 deletion group. There was no significant difference in number of diagnoses between the 22q11.2 deletion group and the other ND-CNV group (t(33.7) = 0.14, *P* = 0.89).

IQ tests were administered using the Wechsler Adult Intelligence Scale (WAIS-III). We report Verbal IQ (VIQ), Performance IQ (PIQ) and Full Scale IQ scores (FSIQ). There was no significant difference in IQ between the 22q11.2 deletion group and the other ND-CNV group (t(22.7-25.1) < 0.57, *P* > 0.58).

In the ND-CNV group, 62% of participants were taking medication for physical, neurological or mood disorders (e.g. high blood pressure, asthma, pain/migraine, depression/anxiety), with the most common medications including gabapentin, co-codamol (combination of codeine and paracetamol), and fluoxetine (N = 3). Given the high variability of medications, their effects could not be systematically investigated; however, their impact was alleviated in MEG analysis by tests of generalizibility (e.g. resampling).

Controls were selected among resting-state datasets acquired at CUBRIC as part of the “100 Brains”’ and “UK MEG Partnership” projects. These cohorts included healthy participants aged 18-65 with no history of neurological or neuropsychiatric disorders, and 42 controls were chosen to match the gender and age of the ND-CNV carriers as closely as possible (22 female; mean age 33.3±9.6 years). These measurements were acquired under protocols approved by the Cardiff University School of Psychology Ethics Committee.

Since a third of the ND-CNV cohort consisted of participants with 22q11.2 deletions, we assessed the impact of this subgroup by repeating all analyses on (1) participants with other ND-CNVs (except 22q11.2 deletions) and their matched controls (N = 56), and (2) participants with 22q11.2 deletions and their matched controls (N = 28).

### Genotyping

Participants with ND-CNVs were genotyped using the Illumina HumanCoreExome whole genome SNP array, which contained an additional 27 000 genetic variants at loci previously linked to neurodevelopmental disorders, including CNVs. The raw intensity data was processed using Illumina Genome Studio software (version 2011.1). PennCNV (version 1.0.3) was used to perform CNV calling in order to confirm the presence of the ND-CNV in each case sample, with each CNV being required to span a minimum of 10 informative SNPs and to be at least 10 kb in length. CNV coordinates were specified according to genome version hg19, and the boundaries of each CNV were confirmed by manually inspecting the Log R ratio and B allele frequency plots at each of the genomic regions of interest (Table 1). Genetic information was not available for control participants; given the rarity of these genotypes in the general population, they were assumed to carry no ND-CNVs.

### Data collection

Five-minute resting-state MEG recordings were made using a 275-channel CTF radial gradiometer system (CTF, Vancouver, Canada) at a sampling rate of 1200 Hz. Three of the sensors were turned off due to excessive noise, and 29 reference channels were recorded to improve noise cancellation(Vrba, 2002). During the recordings, participants were seated upright and fixated their eyes on a red fixation point presented centrally on either a CRT monitor or LCD projector. Three electromagnetic coils were placed at fiducial locations (nasion and pre-auricular) for head localization.

To aid in source localization, structural T1-weighted MRI scans were also acquired using a 3T General Electric or Siemens MRI scanner with a 1 mm isotropic FSPGR/MPRAGE pulse sequence.

### Data analysis

#### Pre-processing

To remove muscle artifacts, a semi-automatic procedure was implemented using the FieldTrip toolbox (Oostenveld et al., 2011) and MATLAB R2015a. Sensor time-series were bandpass-filtered between 110 and 140 Hz and z-transformed; segments exceeding a participant-specific z-score threshold were removed. Next, eye movement and cardiac artifacts were projected out of the data using Independent Component Analysis (ICA). Noisy channels exhibiting high variance were also removed from the data where necessary. There was no significant difference in recording duration after artifact rejection between the ND-CNV and control groups (t(81.8) = 1.61, *P* = 0.11, mean duration 255.88±29.31 s and 245.33±30.86 s respectively).

Head motion was monitored continuously in 18/42 ND-CNV datasets and 40/42 control datasets, and head localization was performed at the start and end of the recording in the remaining datasets. There was no significant difference between the ND-CNV and control groups in maximum head coil displacement between the beginning and end of the recording (t(79.8) = 0.85, *P* = 0.39, mean displacement 2.07±3.62 mm and 2.7±3.06 mm respectively). In datasets with continuous head localization, the maximum distance of the head coils from their average position across the entire recording did not significantly differ between groups (t(39.4) = 1.44, *P* = 0.16, mean distance 4.74±3.5 mm and 3.21±4.2 mm respectively).

Prior to source localization, coregistration was performed by manually marking head coil locations on each participant’s MRI using FieldTrip. The data were downsampled to 600 Hz and bandpass-filtered in six different frequency bands: delta (2-4 Hz), theta (4-8 Hz), alpha (8-13 Hz), beta (13-30 Hz), low gamma (40-60 Hz), and high gamma (60-90 Hz).

#### Estimating functional connectivity

To assess group differences in resting-state connectivity (Figure 1), we focused on amplitude-amplitude coupling of source-localized oscillatory signals (Colclough et al., 2016). Continuous data in each of the six frequency bands were projected into source space using a Linearly Constrained Minimum Variance (LCMV) beamformer. Sources were reconstructed on a 6 mm template grid warped to each participant’s MRI, using a multiple local-spheres forward model (Huang et al., 1999). To alleviate the depth bias, beamformer weights were normalized by their vector norm (Hillebrand et al., 2012).

**Figure 1.**
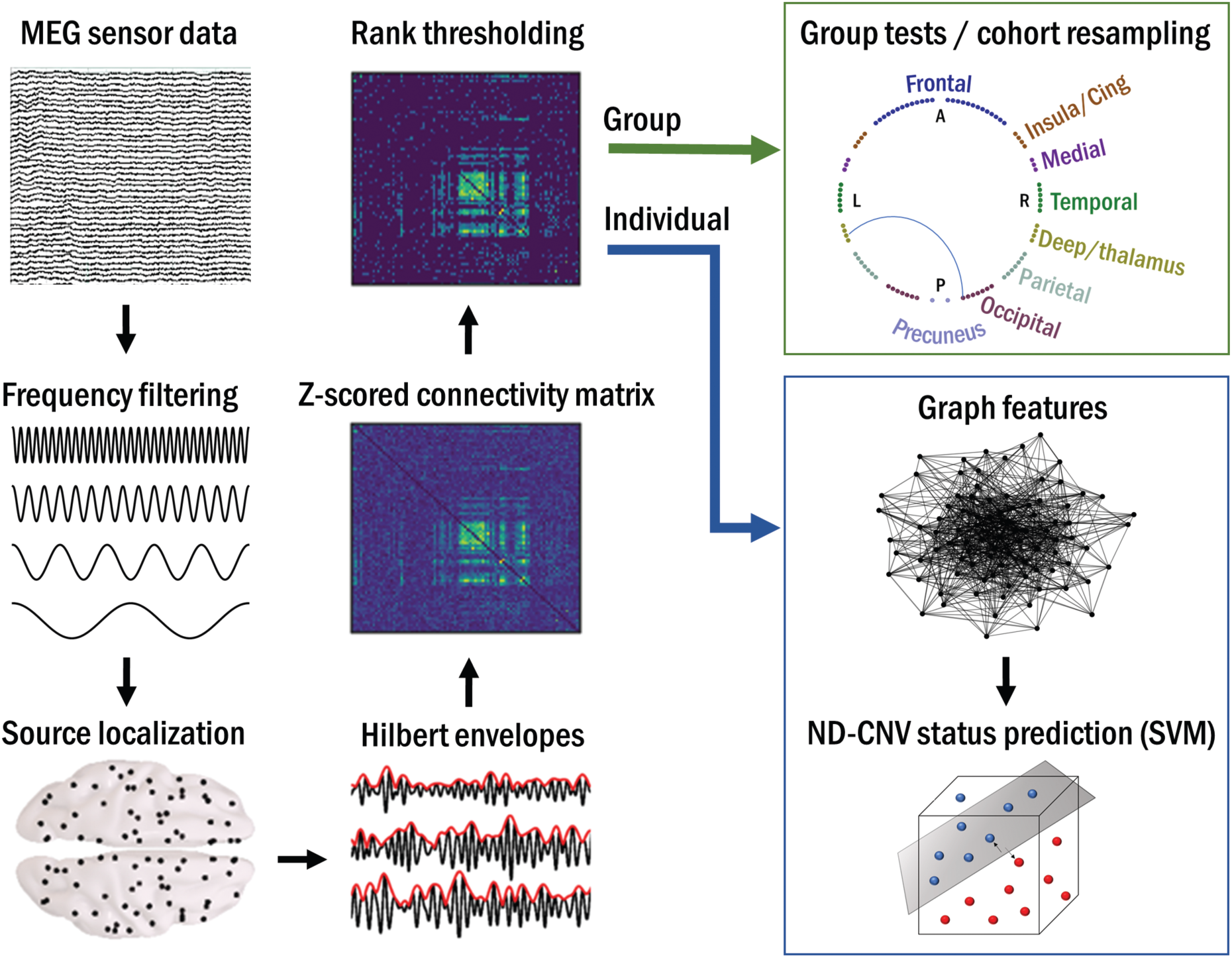
Overview of the analysis pipeline. Resting-state MEG data were preprocessed, filtered into six frequency bands, and projected into source space. Hilbert envelopes were calculated at 90 AAL-atlas-based virtual sensor locations, and correlated to obtain functional connectivity matrices. These were z-scored and rank-thresholded at the group level for between-group analyses, and at the subject level for data-driven prediction of ND-CNV status using graph theory.

Next, 90 nodes corresponding to cortical regions of interest (ROI) in the Automated Anatomical Labelling (AAL) atlas (Tzourio-Mazoyer et al., 2002) were identified by performing a frequency analysis on all sources within each ROI and selecting the source with the largest temporal standard deviation. Continuous virtual sensor timecourses corresponding to the 90 nodes were then reconstructed and bandpass-filtered into the frequency bands of interest.

To avoid spurious correlations, the node time-series were orthogonalized using a multivariate symmetric orthogonalization approach (Colclough et al., 2015). A Hilbert transform was used to calculate oscillatory amplitude envelopes, which were then despiked using a median filter, downsampled to 1 Hz, and trimmed to avoid filter and edge effects. To obtain connectivity matrices, pairwise correlations were calculated between the 90 Hilbert envelopes. Next, a Fisher transform was applied to obtain z-scores with zero mean and unit variance across connections in each participant’s map. This procedure corrected for possible systematic differences across participants, for example due to differences in data quality (Siems et al., 2016).

Intracranial volume (ICV), quantified as the number of 1-mm isotropic voxels inside the brain, was smaller in the ND-CNV group than the control group (t(65.5) = -2.19, *P* = 0.03), in line with some previous reports (Hu et al., 2017; Sønderby et al., 2018; Warland et al., 2019). The potential impact of this difference on the MEG results was alleviated through the source localization procedure and the z-scoring of the connectivity matrices.

In addition to the six frequency bands listed above, a combined measure of connectivity was obtained by calculating the vector-sum of connectivity matrices across all frequency bands (Koelewijn et al., 2019).

#### Group differences in resting-state connectivity

To reduce the impact of noise, a conservative ranking procedure(Koelewijn et al., 2019) was used to threshold the connectivity maps for the purposes of between-group comparisons. This consisted of calculating the rank of each connection in participant-level connectivity matrices and averaging the resulting rank map across participants in each group. Only the top 20% edges in the average rank map were considered “valid” and selected for further analysis. To ensure that large differences in signal across cohorts were not discarded by this procedure, the rank-thresholding procedure was performed separately in each cohort, and connections determined as “valid” in either cohort were included in further analysis.

To assess differences between groups, Welch’s t-tests were conducted at each valid edge. Significant edges were identified using an uncorrected α = 0.05. Correction for multiple comparisons was applied using a randomization procedure with 10000 sign-shuffling iterations and maximal statistic thresholding (omnibus α = 0.05)(Nichols and Holmes, 2001).

The robustness of cohort differences was evaluated through a resampling procedure. Increases and decreases in connectivity between groups were tabulated using random samples of half of each group. This was repeated across 10000 iterations, and edges showing a consistent effect direction across at least 95% of iterations were considered robust.

To control for potential confounds (for example, resulting from imperfect age matching between the ND-CNV and control groups), an additional multiple regression analysis was performed. Combined-frequency connectivity matrices were entered as response variables with a categorical predictor (ND-CNV presence) and three covariates (age, gender, and ICV). A resampling procedure as described above was performed to assess the robustness of between-group differences. The sign of the regression slope associated with the main predictor was tabulated across 1000 split-half cohort randomizations. Edges showing consistent effects across 95% of iterations were considered robust.

#### Individual networks: identifying participants with ND-CNVs using graph theory

Next, a data-driven graph theory approach was used to assess whether participants with ND-CNVs could be distinguished from unaffected controls using functional connectivity features. To this aim, the cohort was divided into training and test sets using an iterated cross-validation procedure.

This analysis focused on individual networks by selecting the top 20% of connections in each participant’s normalized connectivity map as the basis for undirected graphs. Networks were then characterized using six nodal graph theory metrics. These included three measures of node connectedness: *degree* (the number of connections linking each node to other nodes); *betweenness centrality* (the fraction of shortest paths between nodes containing a given node); and *eccentricity* (the maximal shortest path from a node to any other node). *Global efficiency* (the average inverse shortest path between a node and all others) was evaluated as a measure of network integration. Finally, two metrics captured network modularity: *local efficiency* (the average inverse shortest path between a node and its neighbours) and *clustering coefficient* (the fraction of connected node triplets around a node). All metrics with the exception of node degrees were weighted by the inverse of the normalized connectivity matrices; in other words, stronger amplitude-amplitude coupling was taken to reflect shorter paths between nodes. Graph theory analyses were performed using MATLAB R2019a and the Brain Connectivity Toolbox (Rubinov and Sporns, 2010).

To discriminate between groups, a linear support vector machine (SVM) classifier (Boser et al., 1992) was trained on each of the node metrics. Additionally, a pooled feature vector was created by combining the six metrics to maximize the amount of complementary information input to the classifier. This approach makes use of information across all nodes while avoiding the need for multiple testing.

Classification was performed between the ND-CNV and control groups, as well as between the two ND-CNV subgroups (22q11.2 deletions and other ND-CNVs) and their matched controls. To avoid overfitting, model performance was evaluated using 100 iterations of stratified five-fold cross-validation. This entailed iteratively leaving out a fifth of the data for testing and training the model on the remaining data, whilst ensuring balanced group representation in each fold. Performance was quantified using accuracy (proportion correctly classified observations), sensitivity (true positive rate) and specificity (true negative rate) in order to highlight any asymmetries in ND-CNV and control identification. Furthermore, significance was assessed by shuffling the true labels 5000 times and recomputing classifier accuracy to estimate the empirical chance level and calculate a one-tailed p-value (Nichols and Holmes, 2001).

## Results

### Connectivity alterations associated with ND-CNVs

The analysis of group differences in oscillatory connectivity revealed the largest number of valid connections in the alpha and beta bands (Figure 2). Most of the significantly different connections showed a decrease in oscillatory connectivity between posterior, parietal and temporal nodes in the ND-CNV group, with the exception of a few right-hemisphere edges. More extensive cohort effects were detected using the combined frequency maps (61 edges exceeded the uncorrected threshold, compared to 1, 28 and 42 in the theta, alpha and beta bands). These patterns were robust to random sub-sampling of the cohort, suggesting that they were not driven by individual subjects. A small number of left-hemisphere connections, including the precuneus, early visual cortex, and parietal regions, survived omnibus correction for multiple comparisons.

**Figure 2.**
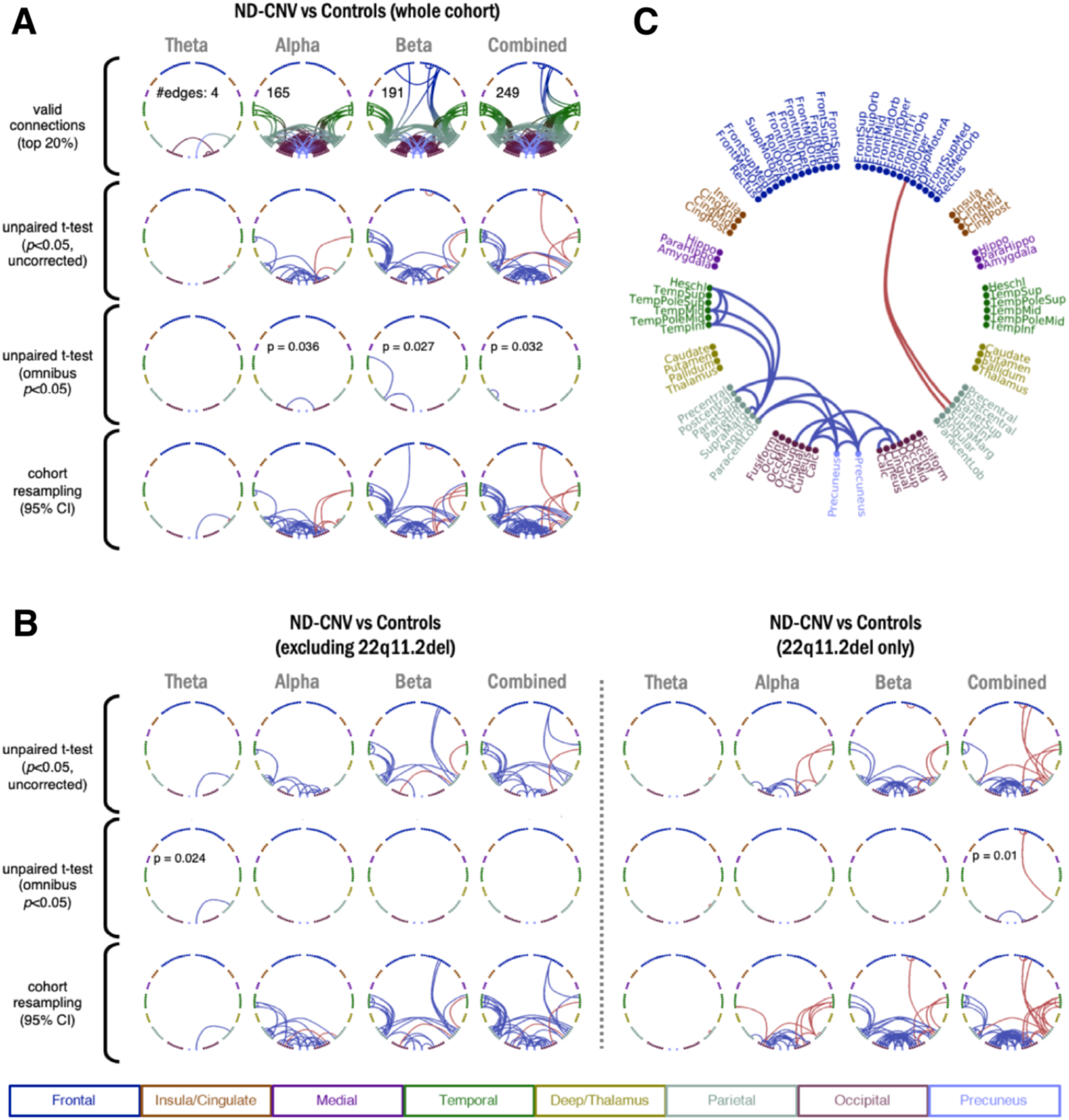
**A.** Differences in resting-state connectivity (amplitude correlations) between participants with ND-CNVs and controls. Connectivity increases and decreases in the ND-CNV group are shown in red and blue respectively. The rows show (top to bottom): valid connections after mean-rank thresholding in each frequency band; uncorrected (*P*<0.05) differences between groups; multiple comparison-corrected (omnibus *P*<0.05) differences between groups; and connections showing consistent increases/decreases in 95% of cohort resampling iterations. **B.** As in A, for subgroups excluding participants with 22q11.2 deletions and their matched controls (left) or including only participants with 22q11.2 deletions and their matched controls (right). To facilitate comparison, “valid” connections were the same as in A. Only frequency bands with surviving “valid” connections are shown. **C.** Nodes displayed in the circular plots, labelled and colour-coded by region. Connections shown are the result of the conjunction analysis between the 22q11.2 deletion group and the other ND-CNV group (also see Figure 3: blue connections are decreased in both groups; red connections have opposing signs).

Importantly, a similar pattern of hypoconnectivity was observed even after excluding participants with 22q11.2 deletions and their matched controls (Figure 2B-C). Both ND-CNV subgroups showed decreased posterior connectivity (Figure 3; 18 connections decreased in both groups), indicating that the overall pattern was not driven by the 22q11.2 deletion group. This pattern occurred despite higher heterogeneity in the mixed ND-CNV group (Figure 4) and might thus reflect convergent alterations across genotypes. On the other hand, participants with 22q11.2 deletions exhibited more right-hemisphere hyperconnectivity compared to controls. These effects spanned the precuneus and parietal cortex, as well as frontal regions, suggesting some overlap with the default mode network (DMN).

**Figure 3.**
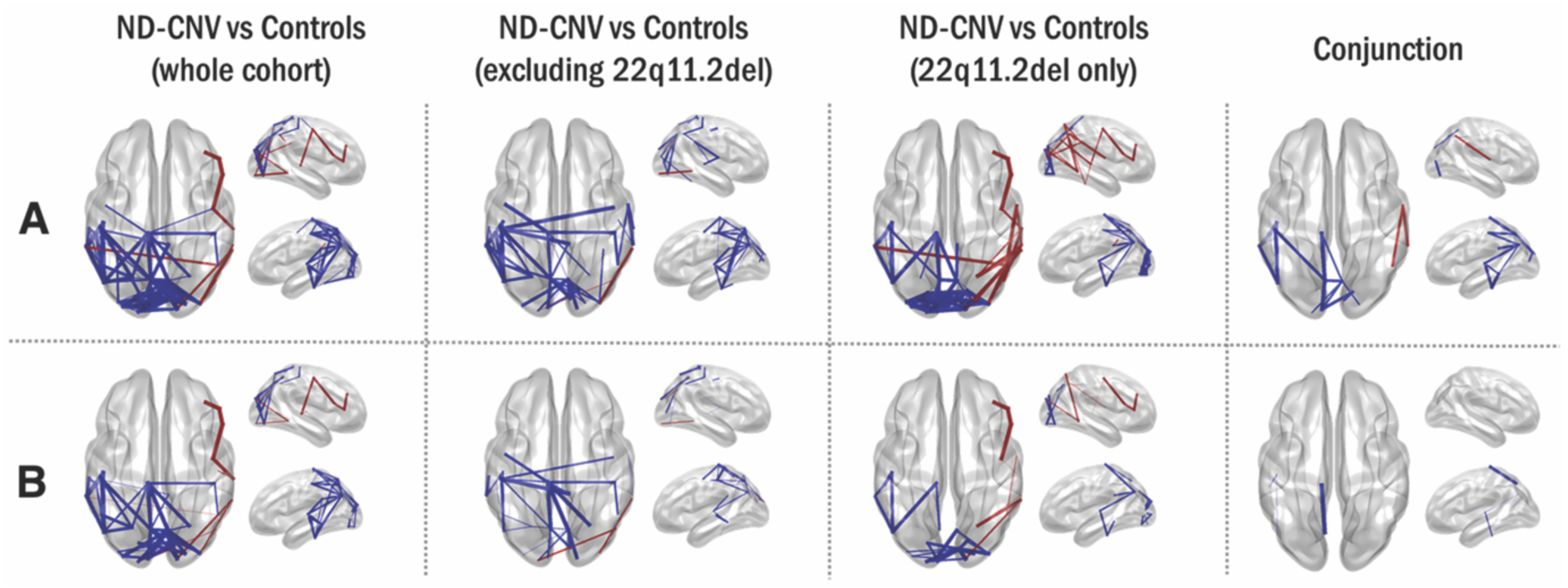
Differences in connectivity are not driven by age, gender, and intracranial volume. Connections meeting the 95% confidence criterion in the cohort resampling test are displayed for all group tests (first three columns). The last column shows supra-threshold connections in both the 22q11.2 deletion group and the other ND-CNV group; they are shown in blue where they are decreased in both groups, and in red where they have opposite signs. Line width increases with effect robustness. **A**. Connections exhibiting robust differences based on the cohort resampling test of combined frequency matrices, plotted on the template brain. **B.** As in A, after including age, gender and intracranial volume as covariates in a multiple linear regression with “ND-CNV presence” as a main categorical predictor.

**Figure 4.**
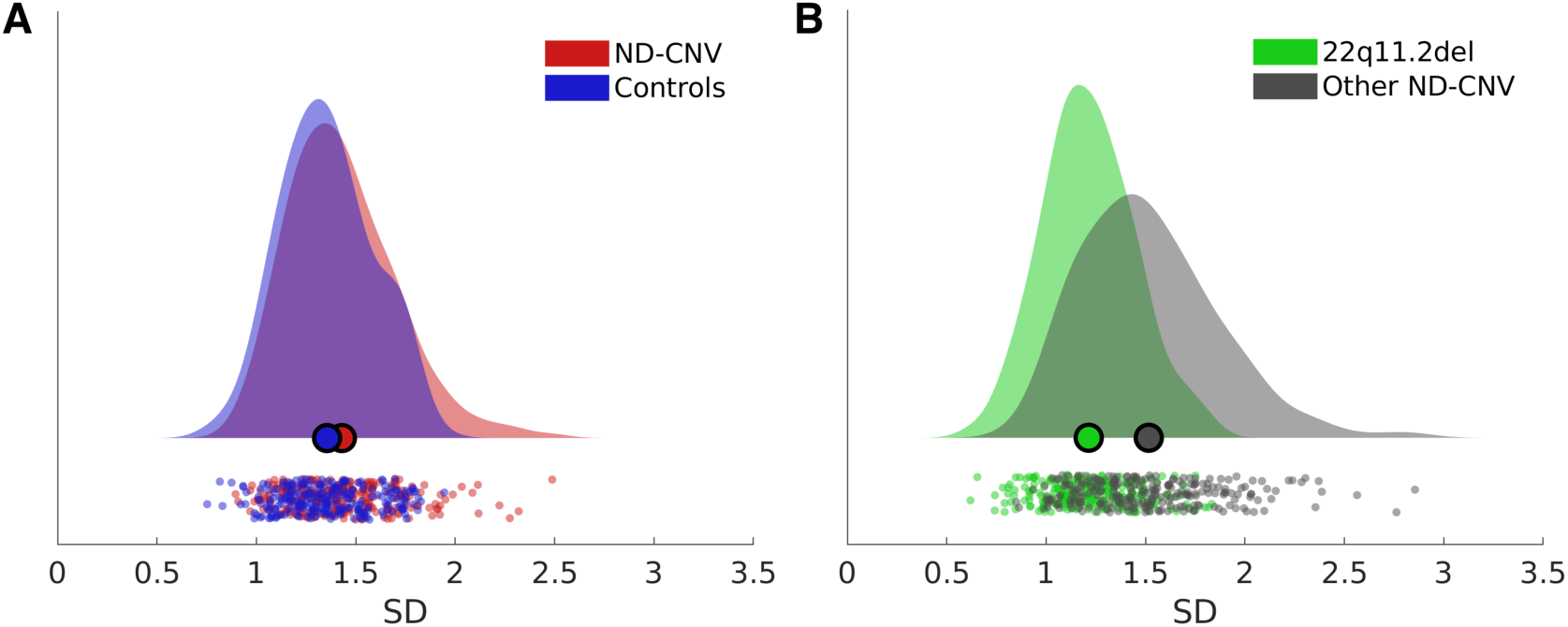
Variability in connection z-scores across participant groups. The distribution of standard deviations (SD) across valid connections in the combined frequency maps are shown for: **A.** the ND-CNV and control groups; **B.** the two ND-CNV subgroups. The mixed ND-CNV group displays larger variability. Plots were generated using the RainCloudPlots toolbox (Allen et al., 2019).

To ensure that these differences were not affected by potential confounds, the cohort resampling tests on combined frequency maps were repeated as a multiple linear regression with age, gender and ICV as covariates. This analysis revealed fewer connections (65 compared to the original 92 in the whole cohort analysis), but largely similar patterns of dysconnectivity (Figure 3B).

Furthermore, although IQ scores could not be included in this analysis because they were not available for the control group, IQ scores in the ND-CNV group significantly correlated with connectivity strength at only four edges (max Pearson’s *r* = 0.5, *P*<0.05).

### Network features as predictors of ND-CNV status

A graph theory framework was employed to identify participants with ND-CNVs from their functional networks based on combined frequency maps. This approach has the advantage of reducing dimensionality and complements the edge-focused group testing approach described above.

Graph theory metrics were successful predictors of ND-CNV participants relative to unaffected controls. The best prediction accuracy was achieved by combining the 6 node features (Table 2; Figure 5A; maximum accuracy 71%±3.44, *P*=0.0002). However, participants with 22q11.2 deletions were more consistently correctly classified than those with other ND-CNVs (Figure 5B).

**Figure 5.**
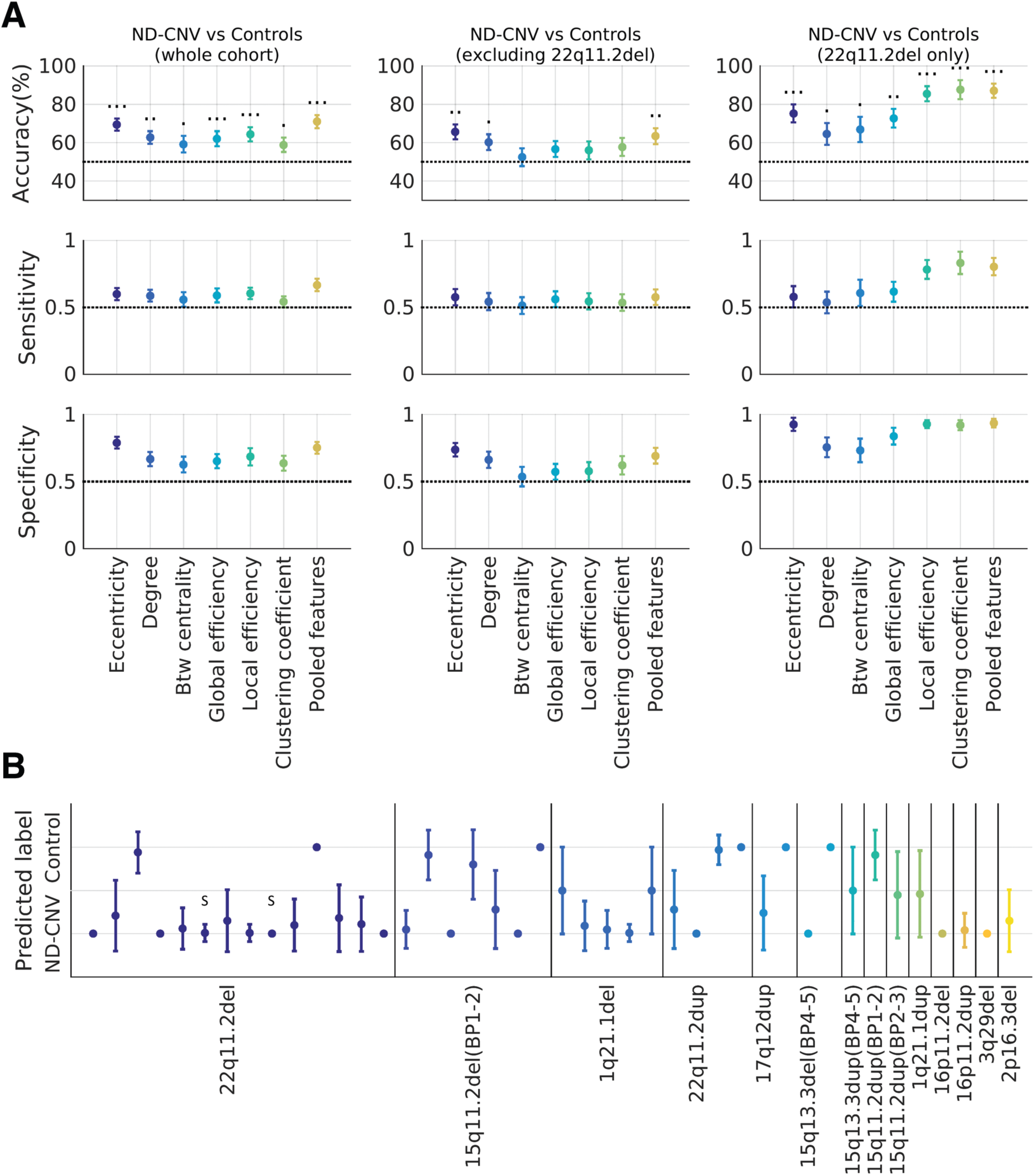
Classifying participants with ND-CNVs and unaffected controls from individual MEG functional networks using graph theory. **A:** Classification performance on the three groups, using different metrics to characterize the networks (eccentricity, degree, betweenness centrality, global and local efficiency, clustering coefficient, and the pooled model). Above-chance accuracies (permutation testing) are marked with 1, 2, and 3 dots respectively for *p*<0.05, *p*<0.01, and *p*<0.001. **B.** How well are different ND-CNVs classified? The plot shows the mean predicted label for each of the 42 participants with ND-CNVs across 100 cross-validation iterations using pooled node features. Participants with 22q11.2 deletions are most consistently correctly identified. Two participants with schizophrenia diagnoses are marked with “S”.

**Table 2.** Network-based classification results. Note that cross-decoding across ND-CNV subgroups achieves high specificity but not sensitivity (middle-right and lower-right cells). This suggests that classifier performance is driven mainly by network features in the control group. For classification between participants with 22q11.2 deletions and other ND-CNVs (top right cell), 100 subsamples of 14 participants were drawn from the 28 carriers of other ND-CNVs, gender-matched to the 22q11.2 deletion group. The subgroups were similar in age, ensuring balanced classes in terms of sample size, gender, and age. Predicted labels were then averaged across cohort resamplings (mean 50±5.57 repetitions per participant).

|  | ND-CNV vs Controls (whole cohort) |  |  |  |  |  |  | 22q11.2del<br>vs other ND-CNV |
| --- | --- | --- | --- | --- | --- | --- | --- | --- |
|  | *Eccentricity* | *Degree* | *Betweenness*<br>*centrality* | *Global*<br>*efficiency* | *Local*<br>*efficiency* | *Clustering*<br>*coefficient* | *Pooled*<br>*node*<br>*features* | *Pooled node*<br>*features* |
| Accuracy (%) | 69.41 | 62.74 | 59.25 | 62.1 | 64.44 | 58.9 | 71 | 72.43 |
| SD | 3.14 | 3.25 | 4.3 | 3.91 | 3.7 | 3.72 | 3.44 | 1.63 |
| P-value | 0.0002 | 0.0012 | 0.0108 | 0.001 | 0.0006 | 0.022 | 0.0002 | 0.0036 |
| Sensitivity | 0.6 | 0.59 | 0.56 | 0.59 | 0.6 | 0.54 | 0.67 | 0.67 |
| Specificity | 0.79 | 0.67 | 0.63 | 0.65 | 0.68 | 0.64 | 0.75 | 0.75 |
| ND-CNV vs Controls (excluding 22q11.2del and matched controls) |  |  |  |  |  |  |  | Train on<br>22q11.2del, test<br>on other ND-CNV |
| Accuracy (%) | 65.62 | 60.27 | 52.45 | 56.69 | 56.06 | 57.82 | 63.43 | 55.36 |
| SD | 3.91 | 4.12 | 4.65 | 4.14 | 4.7 | 4.72 | 4.16 | - |
| P-value | 0.0016 | 0.0242 | 0.2967 | 0.1038 | 0.1032 | 0.053 | 0.0068 | 0.225 |
| Sensitivity | 0.58 | 0.54 | 0.51 | 0.56 | 0.55 | 0.53 | 0.58 | 0.21 |
| Specificity | 0.74 | 0.66 | 0.54 | 0.57 | 0.58 | 0.62 | 0.69 | 0.89 |
| ND-CNV vs Controls (22q11.2del vs matched controls only) |  |  |  |  |  |  |  | Train on other<br>ND-CNV, test<br>on 22q11.2del |
| Accuracy (%) | 75.31 | 64.56 | 66.91 | 72.75 | 85.53 | 87.61 | 87.07 | 75 |
| SD | 4.7 | 5.69 | 6.52 | 4.89 | 3.94 | 4.95 | 3.69 | - |
| P-value | 0.0002 | 0.0296 | 0.0118 | 0.003 | 0.0002 | 0.0002 | 0.0002 | 0.0048 |
| Sensitivity | 0.58 | 0.54 | 0.61 | 0.62 | 0.78 | 0.83 | 0.8 | 0.57 |
| Specificity | 0.93 | 0.75 | 0.73 | 0.84 | 0.93 | 0.92 | 0.94 | 0.93 |

This was confirmed by subgroup classification analyses, which also pointed to subgroup differences. When excluding participants with 22q11.2 deletions, the best decoding accuracies were achieved using node eccentricities (65.62%±3.91, *P*=0.0016), node degrees, and the joint feature model. On the other hand, all node features were successful in discriminating participants with 22q11.2 deletions from their matched controls, with the best performance obtained using the clustering coefficient (87.61%±4.95, *P*=0.0002).

These results point to commonalities in network features (such as decreased centrality) that allow for the successful classification of participants with ND-CNVs across distinct genotypes. On the other hand, the features are specific enough to allow successful discrimination between participants with 22q11.2 deletions and other ND-CNVs (Table 2). Given the higher overall burden of 22q11.2 deletions in neurodevelopmental disorders (Kirov et al., 2014), this suggests that increased neurodevelopmental risk may be associated with more salient alterations in network function and may underpin specific genotype effects.

## Discussion

To our knowledge, the present study provides the first insight into oscillatory connectivity alterations in people with rare ND-CNVs. Using both an established group analysis pipeline and a data-driven graph theory framework, we uncovered a pattern of functional dysconnectivity affecting posterior regions in participants with ND-CNVs. These patterns were robust to effects of age, gender, and intracranial volume, and emerged despite a conservative thresholding approach restricted to the most reproducible connections.

Effects originated in the alpha and beta frequency bands, which are thought to underpin long-range communication between brain areas (Schnitzler and Gross, 2005). Connections linking parietal, temporal, and occipital areas were most consistently affected in both the 22q11.2 deletion group and the other ND-CNV group. Similar patterns have been previously reported in schizophrenia patients (Brookes et al., 2016), including alpha-band parietal hypoconnectivity in first-episode schizophrenia (Phalen et al., 2019). Furthermore, posterior structural network alterations have been identified as an early marker of ASD (Lewis et al., 2014), suggesting a link between such alterations and increased neurodevelopmental risk.

Similar connectivity changes in the visual processing system and the default mode network have been shown in people with 22q11.2 deletions using structural and functional MRI (Schreiner et al., 2014; Scariati et al., 2016a; Larsen et al., 2019). Here, we found that these effects are not restricted to the 22q11.2 deletion group, suggesting that long-range connectivity could act as a common marker across genetic variants. Although non-invasive measurements cannot provide direct mechanistic insight, this is consistent with potential alterations in excitatory-inhibitory balance (Deco et al., 2014; Gu et al., 2019) as a mechanism for pleiotropic genetic effects underlying neurodevelopmental disorders (Gao and Penzes, 2015; Foss-Feig et al., 2017; Lee et al., 2017). This is thought to occur through increased excitation or disinhibition caused by gene haploinsufficiency and mediated by impaired GABA and NMDA receptor function (Kehrer et al., 2008; Ramamoorthi and Lin, 2011; Pocklington et al., 2015).

Despite sample size limitations, differences between the two subgroups also point to effects specific to the highly penetrant 22q11.2 deletions. Hypoconnectivity was more extensive in people with other ND-CNVs, while the 22q11.2 deletion group exhibited more focused patterns; these were robust to cohort resampling, suggesting that they are unlikely to be driven by individual cases. These differences were reflected in the graph theory analysis. Although all network features were altered in the 22q11.2 deletion group, their increased modularity was particularly discriminative, in line with previous reports of increased structural network segregation in people with 22q11.2 deletions (Ottet et al., 2013; Scariati et al., 2016b; Sandini et al., 2018). For other ND-CNVs, the only predictive features were centrality measures (specifically the node eccentricity and degree), reflecting hypoconnectivity in the ND-CNV cohort. Between and within-group classification results (Table 2) highlight the ability of graph theory metrics to capture both convergent and specific network alterations, which could help elucidate the link between CNV pathogenicity and neural system function.

We alleviated concerns of potential systematic between-group differences unrelated to genotype by using conservative approaches (e.g. rejecting weaker connections that may introduce noise) and resampling procedures. Although head motion did not appear to differ between groups, this was not continuously measured in all participants and so we could only partially rule out its effects.

Although the present study was able to evaluate ND-CNV effects independently of the contribution of highly penetrant 22q11.2 deletions, the limited sample size remains a concern common in CNV imaging research. The high genotype variability within the cohort makes the specificity of these effects difficult to assess, particularly with regard to differences between the 22q11.2 deletion group and other ND-CNVs. Variability within the heterogeneous ND-CNV group was higher than within the 22q11.2 deletion group (Figure 4), suggesting that our focus on convergent alterations may obscure specific effects. However, the fact that we see differences robust to resampling in this group, despite its heterogeneity, points to common connectivity alterations across distinct genotypes. Studies recruiting larger samples of participants with ND-CNVs, for example through large multi-site collaborations, are necessary to evaluate the generalizability of connectivity patterns and graph theory metrics as “fingerprints” associated with ND-CNV status.

In summary, the present study assessed oscillatory long-range connectivity as a potential marker of pathogenic genetic effects across a range of rare ND-CNVs. Occipital, parietal and temporal brain areas were characterized by consistent hypoconnectivity across genotypes, which was not exclusively driven by the presence of a large number of participants with highly-penetrant 22q11.2 deletions. Functional networks in the ND-CNV group exhibited decreased node centrality and alterations in network efficiency and structure. Furthermore, features specific to highly penetrant variants were present alongside convergent network alterations and enabled successful ND-CNV classification. These results are consistent with a common mechanism for genetic risk, based on an altered balance between excitatory and inhibitory synaptic processes and leading to network dysfunction. We propose that these functional connectivity alterations are an intermediate phenotype on the pathway from synaptic molecular changes to disruption of cognitive function and psychotic illness.

## Acknowledgements

The authors wish to thank: Ffion Evans and Kali Barawi for assistance with psychometric and clinical data collection; Eirini Messaritaki for helpful comments on the graph theory analysis; George Kirov for help with genetic data; and Alexander Shaw for the SourceMesh visualization toolbox. This work was supported by a Wellcome Trust Strategic Award (100202/Z/12/Z), the MRC UK MEG Partnership Grant (MR/K005464/1), and the MRC Doctoral Training Grant (MR/K501086/1).

